# Transcriptional profiles of a foliar fungal endophyte (*Pestalotiopsis*, Ascomycota) and its endohyphal bacterium (*Luteibacter*, Gammaproteobacteria) in co-culture support sulfur exchange and growth regulation

**DOI:** 10.1101/2021.11.24.469969

**Authors:** Justin P. Shaffer, Morgan E. Carter, Joseph E. Spraker, Meara Clark, Brian A. Smith, Kevin L. Hockett, David A. Baltrus, A. Elizabeth Arnold

**Author notes:** Correspondence: Dr. A. Elizabeth Arnold 303 Forbes Building 1140 E. South Campus Drive Tucson, AZ 85721. Co-first authors (Justin P. Shaffer, Morgan E. Carter).

## Abstract

Symbiosis with bacteria is widespread among eukaryotes, including fungi. Bacteria that live within fungal mycelia (endohyphal bacteria) occur in many plant-associated fungi, including diverse Mucoromycota and Dikarya. *Pestalotiopsis* sp. 9143 is a filamentous ascomycete isolated originally as a foliar endophyte of *Platycladus orientalis* (Cupressaceae). It is infected naturally with the endohyphal bacterium *Luteibacter* sp. 9143, which influences auxin and enzyme production by its fungal host. Previous studies have used transcriptomics to examine similar symbioses between endohyphal bacteria and root-associated fungi such as arbuscular mycorrhizal fungi and plant pathogens. However, currently there are no gene expression studies of endohyphal bacteria of Ascomycota, the most species-rich fungal phylum. We developed methods for assessing gene expression by *Pestalotiopsis* sp. and *Luteibacter* sp. when grown in co-culture and when each was grown axenically. Our assays showed that the density of *Luteibacter* sp. in co-culture was greater than in axenic culture, but the opposite was true for the *Pestalotiopsis* sp. Dual RNA-seq data demonstrate that growing in co-culture modulates developmental and metabolic processes in both the fungus and bacterium, potentially through changes in the balance of organic sulfur via methionine acquisition. Our analyses also suggest an unexpected, potential role of the bacterial type VI secretion system in symbiosis establishment, expanding current understanding of the scope and dynamics of fungal-bacterial symbioses.

**TWEET:** When in co-culture, *Luteibacter* downregulates motility and upregulates a T6SS. Gene expression changes in its host, *Pestalotiopsis*, suggest the bacterium impacts fungal cell structure and methionine availability.

**IMPORTANCE:** Interactions between microbes and their hosts have important outcomes for host- and environmental health. Foliar fungal endophytes that infect healthy plants can harbor facultative endosymbionts called endohyphal bacteria, which can influence the outcome of plant-fungus interactions. These bacterial-fungal interactions can be influential but are poorly understood, particularly from a transcriptome perspective. Here, we report on a comparative, dual RNA-seq study examining the gene expression patterns of a foliar fungal endophyte and a facultative endohyphal bacterium when cultured together vs. separately. Our findings support a role for the fungus in providing organic sulfur to the bacterium, potentially through methionine acquisition, and potential involvement of a bacterial type VI secretion system in symbiosis establishment.

This work adds to the growing body of literature characterizing endohyphal bacterial-fungal interactions, with a focus on a model facultative bacterial-fungal symbiosis in two species-rich lineages, the Ascomycota and Proteobacteria.

## INTRODUCTION

Symbioses between eukaryotes and bacteria are widespread, with profound impacts ranging from the benefits of the gut microbiome with respect to human health to the cost of plant pathogens on global agriculture (Lynch and Hsiao, 2019; Savary *et al*., 2019). The molecular mechanisms underlying relationships ranging from antagonism to mutualism have been studied for decades in animals and plants, including the ways in which pathogenic and beneficial microbes establish in a new host. Although ubiquitous in nature, bacterial-fungal interactions remain relatively poorly understood, despite growing knowledge of their contributions to the emergent properties of microbiomes (Araldi-Brondolo *et al*., 2017; Steffan *et al*., 2020). For example, bacteria living with fungi inhabiting plant roots and leaves can influence fungal phenotypes including growth, reproduction, and pathogenicity, as well as the outcomes of plant-fungus interactions (Bianciotto *et al*. 2004; Partida-Martinez & Hertweck 2005; Partida-Martinez *et al*. 2007a; Lumini *et al*. 2007; Shaffer *et al*.2018).

Despite “bacterium-like organelles” being discovered in arbuscular mycorrhizal fungi (AMF) decades ago (MacDonald *et al*., 1982), only recently have endohyphal bacteria (EHB) been identified living intracellularly in diverse plant-associated fungi. To date, members of the Mucoromycota, Basidiomycota, and Ascomycota have been identified as hosts to EHB, including Proteobacteria, Firmicutes, Tenericutes, Bacteroidetes, and others (Bianciotto *et al*. 2003; Partida-Martinez *et al*. 2007b; Hoffman & Arnold 2010; Desiro *et al*. 2015; Shaffer *et al*. 2016). Thus, both the capacity of bacteria to live within fungal hyphae, and the capacity of diverse fungi to harbor bacterial endosymbionts, are phylogenetically widespread and functionally diverse (Araldi-Brondolo *et al*., 2017; Pawlowska *et al*., 2018). For example, one EHB associated with AMF, *Candidatus* Glomeribacter gigasporarum (Betaproteobacteria), is a vertically transmitted, obligate biotroph with a reduced genome (Bianciotto *et al*., 2003; Bianciotto *et al*., 2004). In contrast, diverse EHB cultured from- or observed in ectomycorrhizal fungi and foliar fungal endophytes appear to be facultatively associated with fungal hosts, with relatively unreduced genomes (Izumi *et al*., 2006; Hoffman and Arnold, 2010; Baltrus *et al*., 2017). Even among these facultative interactions, the impacts on fungal hosts by EHB include alterations in carbon use, growth of germinating spores, degradation of plant cell wall compounds, and sporulation (Lumini *et al*., 2007; Partida-Martinez *et al*., 2007a; Arendt, 2015; Shaffer *et al*., 2017). The metabolic, proteomic, and transcriptomic changes that facilitate these associations, and associated bacterial and fungal phenotypes, are not well known. Changes to fungal traits by the presence of a bacterial symbiont may impact other organisms, such as a plant host through increased virulence (Obasa *et al*., 2017) or plant growth promotion (Hoffman *et al.,* 2013; Guo *et al*., 2017a). Therefore, understanding these bacterial-fungal interactions will expand knowledge of fungal ecology more broadly.

*Mycetohabitans* spp. (formerly *Burkholderia*, Betaproteobacteria) and their host, *Rhizopus microsporus* (Mucoromycotina), represent one emerging model system for EHB based on the ability to independently culture and reintroduce the partners *in vitro*. Metabolic and transcriptomic studies have revealed changes in fungal lipid metabolism underlying their partnership formation, and a lack of reactive oxygen species burst in compatible partners (Lastovetsky *et al*., 2016; Lastovetsky *et al*., 2020). The unique requirement of *Mycetohabitans* spp. for *R. microsporus* sporulation provides a context for probing genes involved in fungal reproduction in a genetically recalcitrant fungal clade (Mondo *et al*., 2017). Essential bacterial genes for symbiosis establishment have been identified in *Mycetohabitans* spp., namely a type II and type III secretion system (T2SS, T3SS) required for invading fungal hyphae (Lackner *et al*., 2011; Moebius *et al*., 2014). The T2SS secretes chitinases critical for bacterial entry, but only one T3SS effector protein has been characterized, and it is not required for the establishment of symbiosis (Moebius *et al*., 2014; Carter *et al*., 2020). Yet, many EHB outside of the Burkholderiales lack one or both the T2SS and T3SS (Araldi-Brondolo *et al*., 2017; Baltrus *et al*., 2017). Indeed, the phylogenetic diversity of both EHB and their fungal hosts suggests bacteria can use many yet undiscovered methods to establish in a given fungus, with a variety of potential outcomes. For example, a lipopeptide produced by *Ralstonia solanacearum,* ralsolamycin, induces chlamydospore formation in- and facilitates invasion of multiple fungi (Spraker *et al*., 2016).

An additional emerging model for EHB is *Luteibacter* sp. 9143 (*Xanthomonadaceae*, Gammaproteobacteria) and its host, the foliar fungal endophyte *Pestalotiopsis* sp. 9143 (Sporocadaceae, Xylariales, Ascomycota) (Hoffman and Arnold, 2010). The *Luteibacter*- *Pestalotiopsis* interaction represents the typical facultative, horizontally-transmitted life modes of EHB found in diverse Dikarya (Araldi-Brondolo *et al*., 2017). Unlike *Mycetohabitans* spp., *Luteibacter* sp. 9143 does not have a T3SS, but does have type I, II, IV, and VI secretion systems (Baltrus *et al*., 2017). *Luteibacter* sp. 9143 has not been observed to be vertically transmitted in its host (i.e., it is not observed readily in conidia). It can be isolated reliably in culture (Arendt *et al*., 2016). *Luteibacter* sp. 9143 increases the ability of *Pestalotiopsis* sp. 9143 to establish as an endophyte in plant hosts (Araldi-Brondolo *et al.,* 2017) and enhances its production of indole-3- acetic acid (Hoffman *et al*., 2013). In addition, *Pestalotiopsis* sp. 9143 harboring *Luteibacter* sp. 9143 exhibits increased cellulase activity, growth on cellulose-enriched growth medium, and degradation of senescent leaf tissue (Araldi-Brondolo *et al*., 2017).

In this study, we contextualized phenotypic observations by using a transcriptomic approach to consider gene expression of *Pestalotiopsis* sp. 9143 and its facultative EHB *Luteibacter* sp. 9143, when grown axenically and in co-culture. We used transcripts from axenic cultures of the fungus to optimize assembly and annotation of its genome, and used that modified assembly to map fungal transcripts in parallel to mapping bacterial transcripts to a previously published genome sequence for *Luteibacter* sp. 9143 (Baltrus *et al*., 2017). Our study was designed to test three main predictions. In previous phenotypic assays, we observed that *Luteibacter* sp. 9143 emerges from its fungal host under conditions of stress, and grows readily in a free-living state provided there is sulfur source besides inorganic sulfate (Arendt, 2015; Arendt *et al*., 2016; Baltrus and Arnold, 2017). Therefore, we predicted that the bacterium would be mildly parasitic on the fungal partner, as illustrated by enhanced bacterial growth but reduced fungal growth in co-culture relative to the axenic state. In such a situation we would expect upregulation of genes associated with bacterial growth in co-culture and changes in metabolic genes of both partners related to molecular exchange of sulfur compounds or other metabolites. Second, we predicted that genes upregulated in co-culture would reflect symbiosis-relevant genes, especially if clustered or relevant to secretion systems or nutrient processing, and coinciding with downregulation of genes associated with motility. In the fungus we would expect upregulation of cellular repair mechanisms or other responses to infection, and changes to secondary metabolite genes relevant to symbiotic establishment, such as upregulation of putative signaling molecules. Third, we predicted transcriptional changes related to carbohydrates such as cellulose, based on observed differences in cellulase activity of *Pestalotiopsis* sp. 9143 with and without *Luteibacter* (Araldi-Brondolo *et al*., 2017). To address these predictions, we performed RNA-seq and analysis of differential gene expression comparing *Luteibacter* sp. 9143 and *Pestalotiopsis* sp. 9143 grown together in co-coculture vs. separately (axenically). Here, we report on the genome of *Pestalotiopsis* sp. 9143 as well as the results of our analysis of differential gene expression.

## RESULTS

### Report of the genome of *Pestalotiopsis* sp. 9143

We generated and assembled the genome sequence of *Pestalotiopsis* sp. 9143 grown axenically and following antibiotic treatment (GenBank: JAHZSN000000000.1) by hybrid assembly of Illumina and Oxford Nanopore reads (see *Methods*). The final genome assembly was 46.3 mega base pairs (Mb) with 13,076 predicted protein coding genes and 247 tRNAs (**Table 1**). The BUSCO score, an assessment of genome completeness, was 94.6% representing 1,255 complete genes out of 1,312 (**Table S1**) in the Dikarya subkingdom data set (Seppey *et al*., 2019). Profiling of secondary metabolite clusters using antiSMASH (Blin *et al*., 2019) revealed 64 secondary metabolite regions, four of which included neighboring protoclusters (**Table S2**).

**TABLE 1.** Final statistics for the genome of Pestalotiopsis sp. 9143.

|  |  |
| --- | --- |
| **Genome assembly:** |  |
| Unique scaffolds | 104 |
| Final assembly size | 46,255,514 bp |
| Masked repeats | 1,379,150 bp (2.98%) |
| N50 | 817,980 |
| GC content | 52% |
| **Gene models:** |  |
| Protein-coding | 13,076 chromosomal; 40 mitochondrial |
| tRNAs | 201 chromosomal; 65 mitochondrial |

**TABLE 2.** Differentially expressed genes of *Pestalotiopsis* sp. 9143. annotated with the enriched GO terms associated with the cytoskeleton and cell wall and their top hit using PSI- BLAST.

| Gene ID | Annotation | L2FC | FDR | PSI-BLAST | Organism |
| --- | --- | --- | --- | --- | --- |
| KJ359_001200 | hypothetical protein | 4.42 | 1.83E-05 | Endochitinase | *Manduca sexta* |
| KJ359_008820 | hydrolase 76 protein | 2.94 | 2.54E-04 | Mannan endo-1,6-alpha-mannosidase | *Saccharomyces cerevisiae* |
| KJ359_008681 | hypothetical protein | 2.68 | 1.73E-04 | Kinesin | *Cylindrotheca fusiformis* |
| KJ359_002383 | hypothetical protein | 2.62 | 9.43E-05 | Chitotriosidase-1 | *Homo sapiens* |
| KJ359_012237 | hydrolase 76 protein | 1.67 | 7.40E-04 | Mannan endo-1,6-alpha-mannosidase | *Saccharomyces cerevisiae* |
| KJ359_000369 | Tubulin beta chain (Beta tubulin) | 1.38 | 4.20E-04 | Tubulin beta chain | *Pestalotiopsis microspora* |
| KJ359_011436 | hypothetical protein | 1.32 | 8.61E-03 | Flocculation protein | *Saccharomyces cerevisiae* |
| KJ359_012899 | Chitin synthase, class 2 | 1.30 | 1.78E-03 | Chitin synthase 1 | *Neurospora crassa* |
| KJ359_009573 | hypothetical protein | 1.28 | 8.75E-05 | Chitin Synthase 6 | *Ustilago maydis* |
| KJ359_000339 | hypothetical protein | 1.17 | 2.32E-03 | Proline-rich protein | *Saccharomyces cerevisiae* |
| KJ359_003752 | Chitin synthase, class 1 | 1.16 | 5.09E-03 | Chitin synthase 3 | *Neurospora crassa* |
| KJ359_011288 | alpha-tubulin | 1.12 | 1.65E-03 | Tubulin alpha-B chain | *Neurospora crassa* |
| KJ359_012862 | class II myosin | 1.11 | 8.65E-04 | Myosin 1 | *Magnaporthe oryzae* |
| KJ359_010970 | Myosin type-2 heavy chain 1 | 1.11 | 6.37E-03 | Myosin 2 | *Lachancea kluyveri* |
| KJ359_010970 | Myosin type-2 heavy chain 1 | 1.11 | 6.37E-03 | Myosin 2 | *Saccharomyces cerevisiae* |
| KJ359_005696 | hypothetical protein | 1.06 | 1.11E-03 | Actin-binding protein | *Saccharomyces exiguus* |
| KJ359_012986 | hypothetical protein | -3.06 | 1.93E-13 | Mannan endo-1,6-alpha-mannosidase | *Saccharomyces cerevisiae* |

### Bacterial density and total biomass of experimental cultures supports mild parasitism of *Luteibacter*on *Pestalotiopsis*

Prior to extracting RNA for analysis of differential gene expression, we examined the growth of *Luteibacter* sp. 9143 and *Pestalotiopsis* sp. 9143 grown together in co-coculture as well as each grown axenically. The bacterium was not observed in the axenic fungal culture, nor was the fungus present in the axenic bacterial culture. The average density of free-living *Luteibacter* sp. 9143 (*i.e.*, cells not associated with fungal mycelium) in co-culture flasks was significantly greater than that in axenic culture flasks (Welch *t*-test, *t* = 6.7, *df* = 2.2, *p*-value = 0.02) (**Fig. 1A**). The average density of free-living *Luteibacter* sp. 9143 in co-culture flasks was 8.67 x 10^5^ ± 2.00 x 10^5^ CFU/mL, and that of *Luteibacter* sp. 9143 in axenic culture flasks was 6.67 x 10^4^ ± 4.62 x 10^4^ CFU/mL. The average dried biomass of *Luteibacter* sp. 9143 in axenic bacterial culture flasks was 3.3±0.0 mg. In contrast, we observed significantly reduced growth of *Pestalotiopsis* sp. 9143 in co-culture relative to axenic culture.

**FIG 1.**
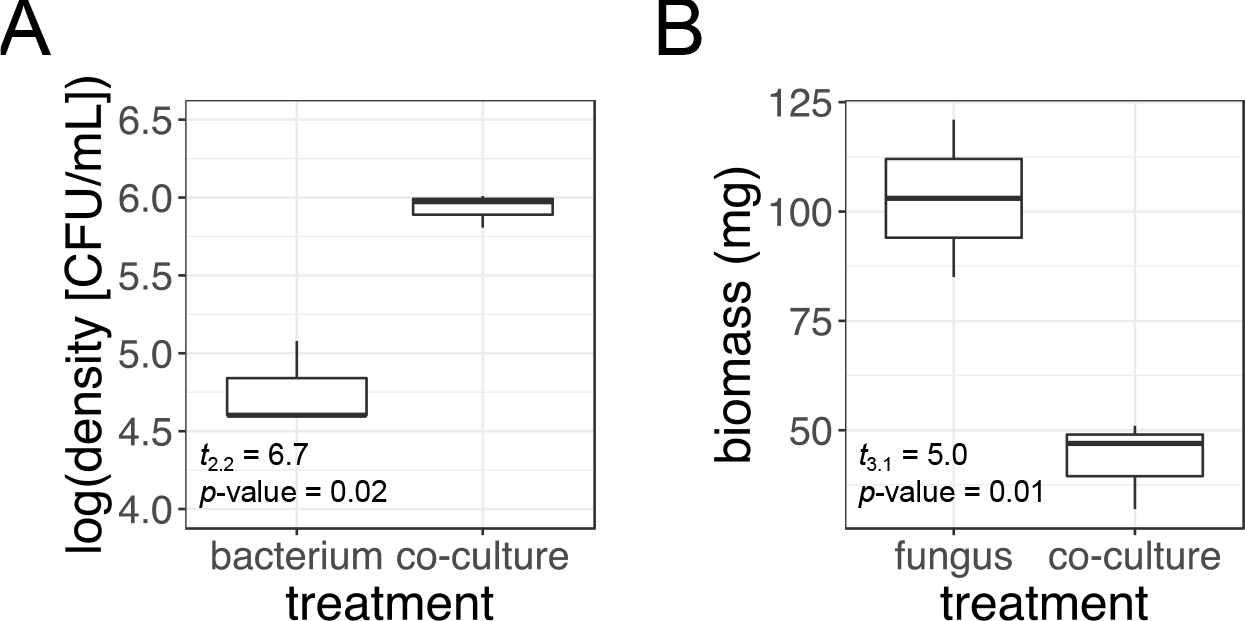
Differences in microbial density/biomass during co-culture vs. axenic growth. (**A**) Density (colony forming units [CFU]/mL) of free-living *Luteibacter* sp. 9143 (*i.e.*, cells not associated with fungal mycelium) when grown axenically (*bacterium*) vs. with *Pestalotiopsis* sp. 9143 (*co-culture*). (**B**) Biomass (mg) of *Pestalotiopsis*sp. 9143 when grown axenically (*fungus*) vs. with *Luteibacter* sp. 9143 (*co-culture*). The same volume of liquid growth media was initially added to each sample. Biomass represents dry mass obtained by filtering cultures to remove the growth medium and lyophilizing the fresh biomass that was retained.

The average combined biomass of *Pestalotiopsis* sp. 9143 and *Luteibacter* sp. 9143 in co-culture flasks was 43.3±0.01 mg, significantly less than 103.00 ± 0.02 mg of *Pestalotiopsis* sp. 9143 in axenic culture flasks (Welch *t*-test, *t* = 5.0, *df* = 3.1, *p*-value = 0.01) (**Fig. 1B**).

### Transcriptional changes in *Luteibacter*sp. 9143 during co-culture reflect increased growth and a symbiotic response

During co-culture with *Pestalotiopsis* sp. 9143, *Luteibacter* sp. 9143 upregulated 217 genes and downregulated 241 genes relative to its gene expression in the axenic state (*i.e.*, for an adjusted *p*-value or false discovery rate [FDR] of 0.01) (**Fig. 2** and **Table S3**). Our analysis of enrichment of gene ontology (GO) terms was largely uninformative, as nearly all terms identified as upregulated were also identified as downregulated. (**Fig. S1**). We observed that 37 of 54 (69%) genes predicted to code for ribosomal proteins were upregulated with a log_2_ fold change (L2FC) of ∼1.5-3.0 (**Table S3**), which we speculate reflects enhanced bacterial growth in co-culture (**Fig. 1A**). Twenty-eight of the 37 (76%) upregulated ribosomal protein genes are encoded in a cluster with four additional ribosomal protein genes not differentially expressed (DE), whereas the remaining nine are scattered throughout the genome (**Table S3**).

**FIG 2.**
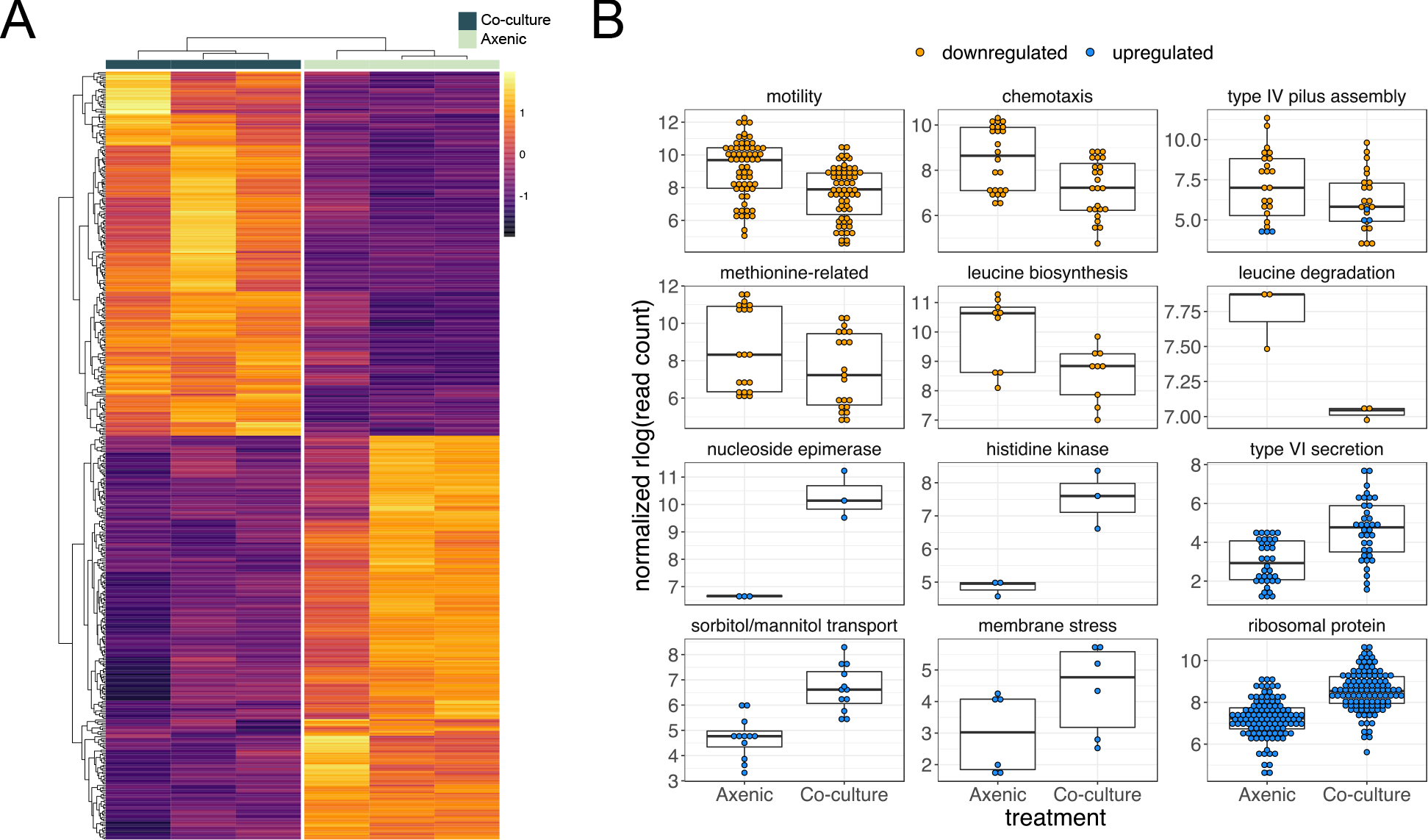
Summary of differentially expressed genes in *Luteibacter* sp. 9143 in co-culture with *Pestalotiopsis* sp. 9143 compared to axenic growth. (**A**) Heatmap showing the log-fold-change (i.e., *deseq2* rlog-normalized) for each gene (rows) for each biological replicate (columns). (**B**) Normalized read counts for axenic vs. co-cultured *Luteibacter* sp. 9143 for specific groups of genes. Methionine-related genes include those for methionine biosynthesis, acquisition, and conversion. Each point represents the read count for a given gene from one replicate culture of a given treatment of *Luteibacter* sp. 9143 (*i.e.*, axenic vs. co-culture), thus three points are present for each gene.

### Luteibacter upregulates type VI secretion, signaling, and transport in co-culture

Genes more strongly upregulated than the majority of the ribosomal proteins included those involved in the type VI secretion system (T6SS). Most known for antibacterial activity, T6SSs serve various roles including metal scavenging, biofilm formation, and interactions with eukaryotic predators and hosts (Bernal *et al*., 2018; Bayer-Santos *et al*., 2019). In total, 11 of 13 (85%) genes predicted to code for T6SS proteins were upregulated, roughly 24% of the 46 genes observed to be significantly and highly upregulated (*i.e.,* adjusted *p*-value ≤ 0.01, L2FC ≥ 3.0; **Fig. 3** and **Table S3**). Similarly, we detected upregulation of two hypothetical proteins within the cluster that are likely co-regulated (**Fig. 3** and **Table S3**).

**FIG 3.**
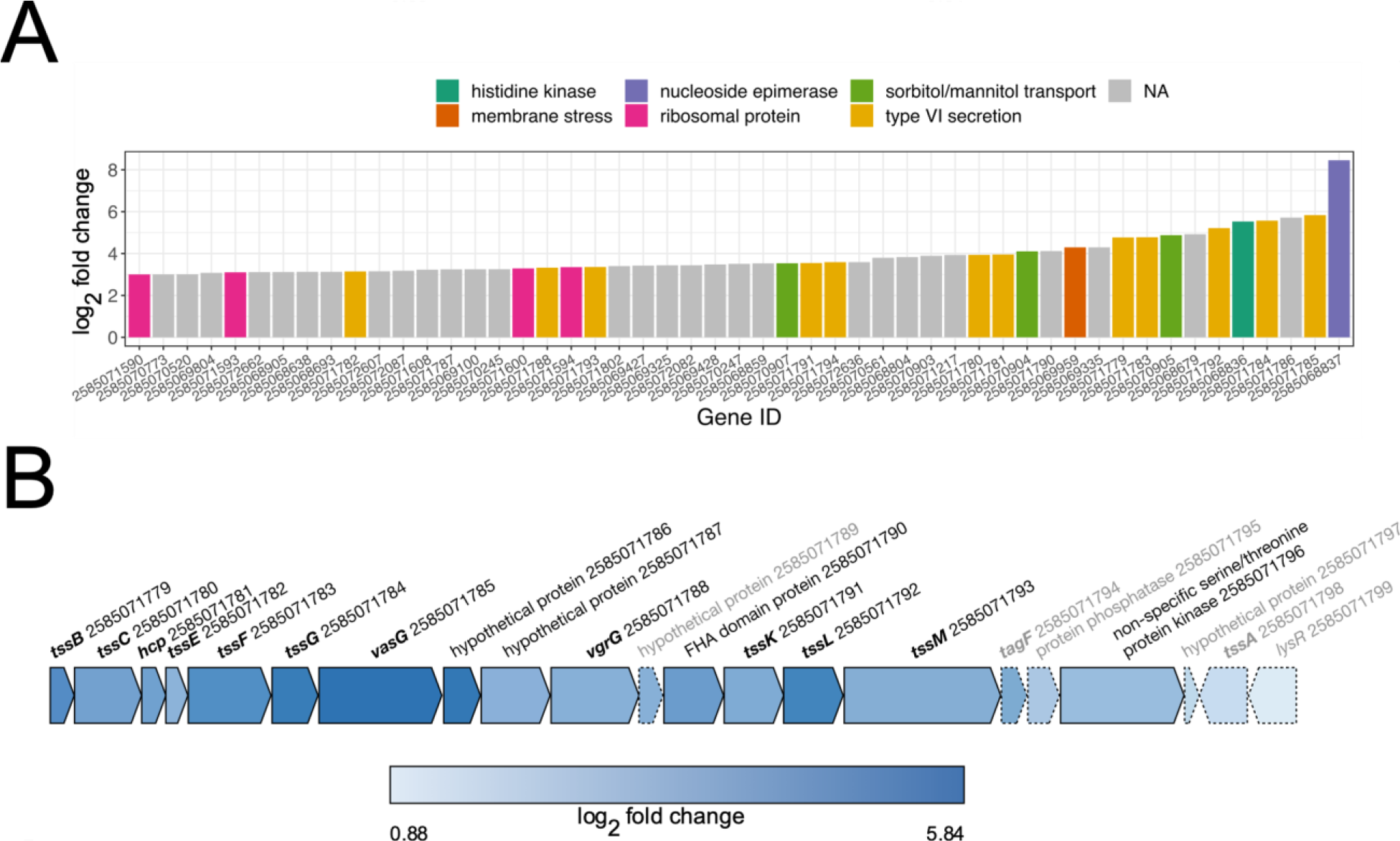
Upregulation of type VI secretion system-related genes in *Luteibacter* sp. 9143 when in co-culture with *Pestalotiopsis* sp. 9143. (**A**) Log_2_ fold change (L2FC) for significantly highly upregulated genes (*i.e.,* adjusted *p*-value ≤ 0.01, L2FC ≥ 3.0), highlighting important groups of genes including those associated with the type VI secretion system (T6SS). (**B**) Physical map showing proximity of genes associated with the T6SS in bold font. Colors indicate L2FC in co-cultured vs. axenic *Luteibacter* sp. 9143. Genes annotated in gray and with hashed lines were neutrally regulated and all others were upregulated.

Additionally, the most- and sixth-most-upregulated genes were also clustered, and were predicted to be a nucleoside-diphosphate-sugar epimerase and a two-component sensor histidine kinase, respectively (**Table S3**). Notably, the response regulator in this cluster is also upregulated although it does not meet our FDR threshold for significance (L2FC = 2.8; *p*-value = 0.02). Two-component regulatory systems composed of kinases and response regulators are key signal transducers for detection of environmental or cellular changes by bacteria.

We also observed all (4/4) genes predicted to be associated with sorbitol/mannitol transport that form a cluster, as well as one adjacent gene predicted to code for a mannitol dehydrogenase, to be upregulated (**Table S3**). Similarly, five of 15 (33%) genes predicted to code for efflux pumps were upregulated.

### Luteibacter downregulates pilus assembly, chemotaxis, and motility in co-culture

We observed many downregulated genes to be associated with type IV pilus assembly, chemotaxis, and motility (**Fig. 2**). Although all 12 genes predicted to code for type IV secretion system (T4SS) proteins were not DE, eight of 24 type IV pilus assembly genes were DE, and nearly all (7/8 or 88%) were downregulated (*i.e.*, *pilA*, *pilB*, *pilC*, *pilQ*, *pilV*, and *pilW*). Only one gene, *pilY1*, was upregulated. For the operon consisting of *pilM*, *pilN*, *pilO*, *pilP*, and *pilQ*, only *pilQ* is downregulated.

Whereas several DE genes predicted to be associated with chemotaxis, flagella, and motility are scattered throughout the genome of *Luteibacter* sp. 9143, the majority were organized in distinct clusters. For example, we found 18 of 41 genes (44%) predicted to code for flagellar proteins and eight of 34 genes (24%) predicted to code for chemotaxis proteins to be downregulated. This included genes predicted to code for the sigma-54 specific transcriptional regulator FliA and an anti-sigma-28 factor in the FlgM family, which are two proteins important for regulating the flagellar protein-coding genes (Osterman *et al*. 2015) (**Table S3**). Along with 15 of the 18 downregulated genes coding for flagellar proteins and three downregulated genes coding for chemotaxis, these genes form a cluster comprising 47 genes related to chemotaxis and motility (**Table S3**).

### Luteibacter downregulates methionine metabolism in co-culture

When considering amino acid metabolism, we observed genes related to methionine to be DE. For example, all (7/7) genes predicted to code for methionine metabolism or transport were downregulated. This included a methionine aminotransferase, a methionine synthase, a methionine adenosyltransferase, two peptide-methionine S-oxide reductases (R and S), a methionine transport system substrate-binding protein, and a homoserine *O*-acetyletransferase, the latter of which was the most downregulated gene in the experiment (**Table S3**). Furthermore, three of four genes involved in leucine biosynthesis and one gene involved in leucine degradation were also downregulated, including the small and large subunits of 3-isopropylmalate dehydratase, 3- isopropylmalate dehydrogenase, and 2-isopropylmalate synthase, and leucine dehydrogenase (**Fig. 2**). This suggests an undescribed role for leucine or other branched chain amino acids during the symbiosis.

### Transcriptional changes in *Pestalotiopsis*sp. 9143 during co-culture may be facilitated by NmrA-like transcription repressors

During co-culture with *Luteibacter* sp. 9143, *Pestalotiopsis* sp. 9143 upregulated 478 genes and downregulated 521 genes for an FDR of 0.01 (**Fig. 4A** and **Table S4**). In contrast to the bacterial genome, the limited number of gene name annotations in the fungal genome made GO analysis a useful tool to identify changes in the fungal transcriptome. There were 21 GO terms enriched in the upregulated genes, and 22 GO terms enriched in the downregulated genes (**Fig. 4B).** The GO terms with the highest enrichment score among upregulated genes were the molecular function (MF) GO terms for ATPase activity (GO:0016887) and ATPase coupled transmembrane transporter activity (GO:0042626). The top biological processes (BP) GO terms were peptide pheromone export (GO:0000770) and methionine metabolic process (GO:0006555), while the three enriched cellular component (CC) GO terms were myosin complex (GO:0016459), integral component of the membrane (GO:0016021), and microtubule (GO:0005874). The enriched GO terms in the downregulated genes had lower enrichment scores overall, with the largest ones being vitamin B6 biosynthetic process (GO:0042819; BP), membrane (GO:0016020; CC), flavin adenine dinucleotide binding (GO:0050660; MF), and oxidoreductase activity, acting on CH−OH group of donors (GO:0016614; MF).

**FIG 4.**
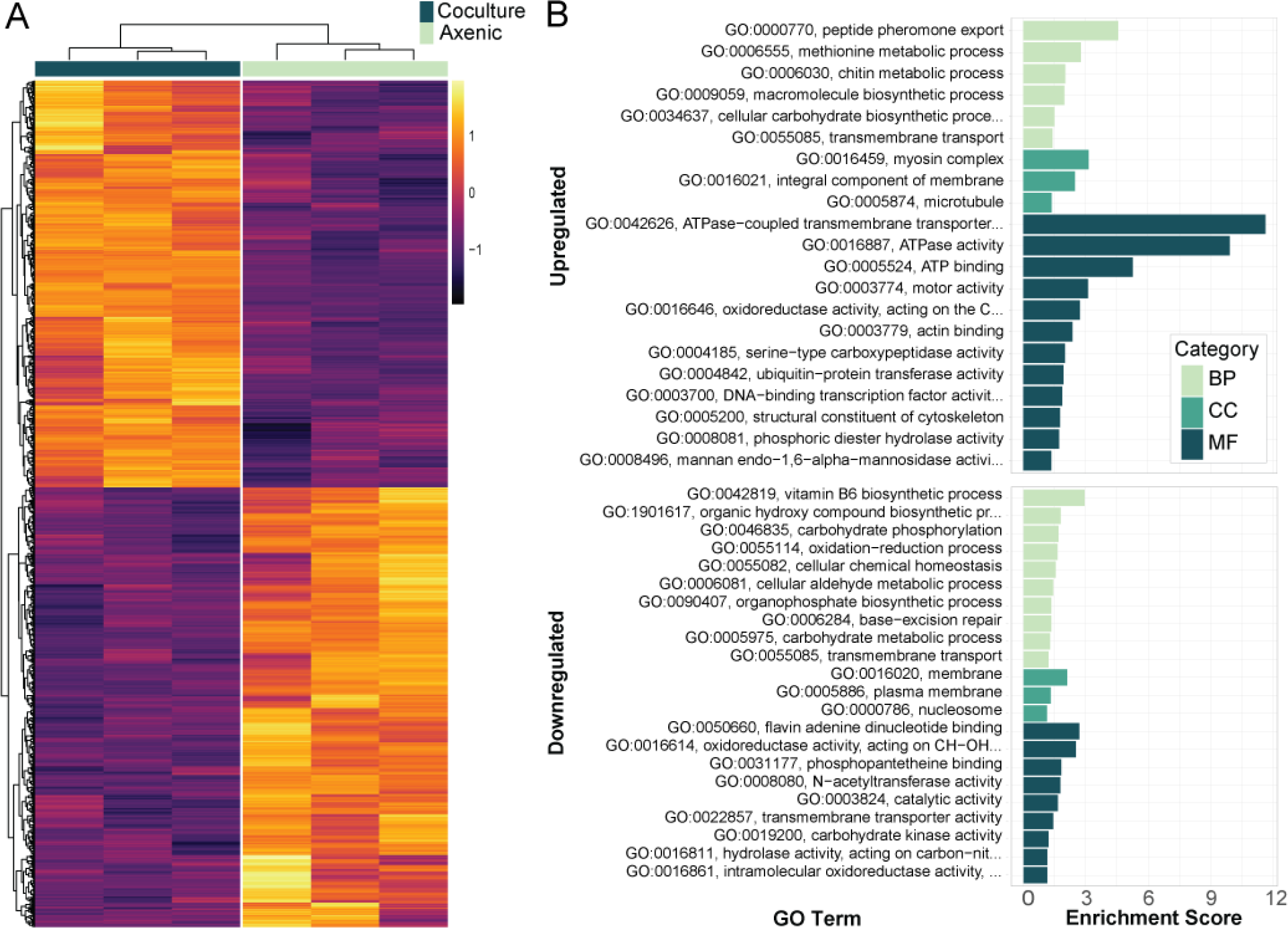
Differentially expressed genes and enriched GO terms in *Pestalotiopsis* sp. 9143 when in co-culture with *Luteibacter* sp. 9143. (**A**) Heatmap of regularized log (rlog) transformed counts showing the differentially expressed genes identified with DESeq2 for a false discovery rate of 0.01. Three biological replicates are shown for co-cultured and axenic fungal growth. (**B**) Enriched Gene Ontology (GO) terms among the up- or downregulated differentially expressed genes in co-cultured fungus; biological processes (BP), cellular component (CC), and molecular function (MF). Enrichment analysis was done using the topGO R package’s weight01 algorithm and Fisher’s exact test for a *p-*value<0.05.

The top two upregulated genes (KJ359_005267 and KJ359_006927) both encode hypothetical proteins with NAD(P)-binding domains that are members of the PFAM NmrA (PF05368), suggesting they are transcription repressors (Andrianopoulos *et al*., 1998). Five additional genes encoding NmrA-like proteins were upregulated, and another one was the ninth most downregulated gene. While a handful of the most upregulated genes are cytochrome P450s and there are some that are downregulated, these are not enriched in the data set as only 14 are differentially expressed out of 234 total. Nearly half of the most downregulated genes (*i.e.*, 13 of 30; L2FC ≥ 6) are hypothetical proteins with no annotated domains of known function. The following sections detail the themes present in the differentially expressed genes and GO term enrichment, highlighting the changes in transporters and cell structure-related genes, methionine and carbohydrate metabolism, and potential defense or signaling compounds like secondary metabolites and β-lactamases.

### Diverse transporters are up- and downregulated by Pestalotiopsis

Transport changes in *Pestalotiopsis* sp. 9143 are considerable in the presence of *Luteibacter* sp. 9143. The term for transmembrane transport (GO:0055085) is enriched in 42 up- and 31downregulated genes that are largely predicted to encode ATP-binding cassette (ABC) transporters or Major Facilitator Superfamily (MFS) transporters, as well as amino acid permeases, oligopeptide transporters, and metal ion transporters. Of the non-MFS and ABC protein-encoding genes, two purine-cytosine permeases (InterPro: IPR001248) are downregulated. Additionally, three genes putatively encoding for magnesium (Mg^2+^) CorA-like transporter proteins are all downregulated, possibly reducing Mg^2+^ ion concentration and contributing to a reduction in growth and cell wall integrity (Reza *et al*., 2016).

Among the downregulated genes associated with transmembrane transport, the majority are MFS transporters (21 of 31) and of those, 11 are from the sugar transporter MFS subfamily (IPR005828 and/or PF00083). In contrast, only five predicted sugar transporters are upregulated including the hexose transporter *hxt1*. Among the genes that were upregulated, 19 ABC transporters contributed to the most enriched GO terms: ATPase-coupled transmembrane transporter (GO:0042646), ATPase activity (GO:0016887), and ATP binding (GO:0005524).

Seven of the 19 are associated with peptide pheromone export (GO:0000770) and only two are predicted to be within secondary metabolite clusters: KJ359_000853 in Cluster 10.1 is upregulated and KJ359_002926 in Cluster 18.2 is downregulated, as mentioned detailed below.

Thus, the substrates of many of these ABC transporters are unknown given the wide variety of potential substrates ranging from lipids to toxins (Perlin *et al*., 2014).

### Cell structure-related gene expression is impacted by bacterial presence

Because manipulation of actin is common in close symbioses, such as rhizobial nodule formation in plant roots (Zhang *et al*., 2019) and intracellular bacterial pathogen motility in animals (Goldberg, 2001), we predicted that cell wall- and membrane-associated genes would be differentially expressed. We found that the upregulated genes are enriched for many GO terms associated with the cytoskeleton and cell wall: chitin metabolic process (GO:0006030), myosin complex (GO:0016459), microtubule (GO:0005874), motor activity (GO:0003774), actin binding (GO:0003779), structural constituent of cytoskeleton (GO:0005200), and mannan endo-1,6- alpha-mannosidase activity (GO:0008496). The genes associated with these terms (**Table 2**) are mostly myosins, chitin synthases, tubulins, and mannan endo-1,6-alpha-mannosidases. Of the three DE mannan endo-1,6-alpha-mannosidases, which are typically required for fungal cell growth and contribute to fungal cell wall biosynthesis (Kitagaki *et al*., 2002), one is actually downregulated. The DE genes of all other enriched GO terms associated with the cytoskeleton or cell wall are upregulated (**Table 2**). These results align with the changes in gene expression seen in other bacterial-fungal symbioses; a compatible *Rhizopus microsporus* isolate upregulates genes involved in cytoskeletal rearrangement and the cell wall when in contact with *Mycetohabitans* spp. (Lastovetsky *et al.,* 2020).

Notably, many of the genes related to cell structure that were not considered differentially expressed for our set FDR<0.01 are just beyond that probability threshold and typically have low L2FCs. For example, of the five genes in the *Pestalotiopsis* sp. 9143 genome annotated with the term myosin complex (GO:0016459), three genes are DE with a L2FC of 1.1-1.2 for an FDR<0.01, but the other two genes are DE for a less stringent FDR<0.05. In filamentous fungi, myosins are associated with proper septation within hyphae, sporulation, and cell wall formation (Guo *et al*., 2017b; Renshaw *et al*., 2016). Additionally, three of the six genes annotated with the MARVEL domain (IPR008253) are downregulated (KJ359_003932, KJ359_001504, and KJ359_002564) for FDR<0.01, whereas two of the remaining three have an FDR<0.06. Proteins containing the MARVEL domain are not well studied in fungi but have been shown to play a role in actin and membrane organization in *Candida albicans* (Douglas *et al*., 2013) and in cell growth and fusion in *Neurospora crassa* (Fu *et al*., 2011).

### Methionine metabolism is upregulated by Pestalotiopsis, complementing downregulation by Luteibacter

Within the upregulated genes, the second most significantly enriched GO term for biological processes is that for methionine metabolic process (GO:0006555) (**Fig. 4B**). Three of 14 genes with this term or child GO terms are slightly upregulated with a L2FC ranging from one to two: two predicted methylenetetrahydrofolate reductases (KJ359_010927 and KJ359_003580) and one methionine synthase (KJ359_000681). Some proteins involved in methionine metabolism use pyridoxal-5’-phosphate (PLP) as a cofactor. However, two genes encoding PLP synthase subunits Snz1 (KJ359_006612) and PdxT/SNO (KJ359_006611) are downregulated, along with a putative PLP oxidase (KJ359_005893). As PLP is a vitamin B6 compound, these genes represent the enrichment of the term for vitamin B6 biosynthetic process (GO:0042819) among the downregulated genes.

### Glycoside hydrolases account for most changes in Pestalotiopsis carbohydrate metabolism during co-culture

We previously observed that cured *Pestalotiopsis* sp. 9143 has reduced cellulase activity and reduced growth on cellulose-based media (Araldi-Brondolo *et al*., 2017), thus we expected to see changes in carbohydrate metabolism when co-cultured. The majority of the downregulated genes (16 of 23) predicted to be involved in the carbohydrate metabolic process (GO:0005975) encode putative members of glycoside hydrolase families; altogether 15 upregulated and 21 downregulated genes are predicted by InterPro to encode glycoside hydrolases. Glycoside hydrolases act on diverse substrates and are commonly secreted out of the fungal cell to break down biomass, including compounds found in plant cell walls like lignin, cellulose, and hemicellulose (Murphy *et al*., 2011). Of the DE glycoside hydrolases, 25 have signal peptides predicted by SignalP (Almagro Armenteros *et al*., 2019).

Two downregulated genes involved in the carbohydrate metabolic process (GO:0005975) are also annotated with carbohydrate phosphorylation (GO:0046835) and encode hexokinases.

The additional carbohydrate metabolism-associated genes are a glucosamine-6-phosphate isomerase, a sugar phosphate isomerase (RpiB/LacA/LacB family), and a polysaccharide deacetylase. The glucosamine-6-phosphate isomerase (KJ359_003080) converts glucosamine-6- phosphate to fructose-6-phosphate and ammonium in the last step of N-acetylglucosamine (GlcNAC) catabolism.

### Only secondary metabolite Cluster 10.1 is fully differentially expressed in co-culture

Of the 64 secondary metabolite clusters (four of which include neighboring clusters) in the *Pestalotiopsis* sp. 9143 genome, 13 have a core biosynthetic gene (i.e., NRPS, PKS, NRPS-PKS hybrid, or terpene cyclase) that is DE when co-cultured with *Luteibacter* sp. 9143 (**Fig. S2**). Six were downregulated and five were upregulated. However, only in Cluster 10.1 are the core PKS genes and neighboring biosynthetic and transport genes all DE (**Fig. 5A, B,** and **Fig. S2**). Cluster 18.2 has a DE core NRPS (KJ359_002925) and ABC transporter gene (KJ359_002926), but they are downregulated and the additional α/β hydrolase (KJ359_002922) predicted to be involved in biosynthesis is not DE (**Fig. 5A, C,** and **Fig. S2**).

**FIG 5.**
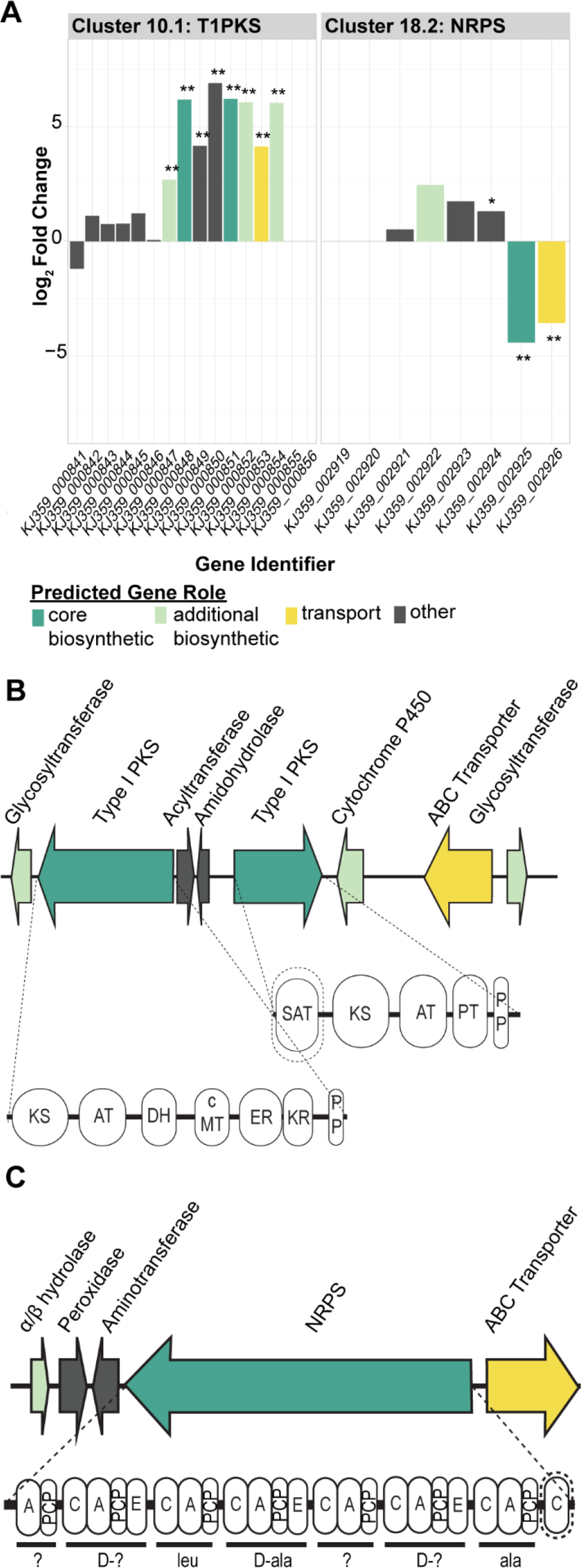
Differential expression of genes within two secondary metabolite clusters of *Pestalotiopsis*sp. 9143. (**A**) Log_2_ fold change of the genes within Cluster 10.1 and Cluster 18.2 color coded by predicted gene role. Two asterisks (**) designate differentially expressed (DE) genes with FDR<0.01; one asterisk (*) designates DE genes with FDR<0.05. (**B**) Diagram of part of the Cluster 10.1 Type I polyketide synthase (PKS) locus with putative gene products, including PKS domains: ketosynthase (KS), acyltransferase (AT), phosphopanetheine (PP), acyl carrier protein (ACP) dehydratase (DH), enoylreductase (ER), ketoreductase (KR), carbon methyltransferase (cMT), product template (PT), and a N-terminal ACP transacylase similar to the one involved in aflatoxin biosynthesis (SAT). (**C**) Diagram of part of the Cluster 18.2 nonribosomal peptide synthase (NRPS) locus with putative gene products, including NRPS domains: condensation (C), AMP-binding (A), epimerization (E), and peptidyl-carrier protein (PCP). Bars underneath groups of ovals indicate NRPS modules and the amino acids they are predicted to load: alanine (ala) and leucine (leu). Incomplete modules or predictions are indicated with a question mark (?).

Cluster 10.1 contains genes putatively encoding two polyketide synthases (PKS), two glycosyltransferases, a cytochrome P450, and an ABC transporter (**Fig. 5B**). Additional genes for an acyltransferase and an amidohydrolase were not predicted as serving a role in the cluster, but are upregulated and may also contribute to a final product. The eight genes in total that were upregulated in Cluster 10.1 ranged from a L2FC of 2.5 to 7. Both predicted PKS proteins encoded by KJ359_00848 and KJ359_00851 have the required ketosynthase (KS), acyltransferase (AT), and phosphopanetheine (PP) acyl carrier protein (ACP) group domains.

KJ359_00848 has a carbon methyltransferase domain (cMT) and three additional reducing domains: dehydratase (DH), enoylreductase (ER), and ketoreductase (KR). In contrast, KJ359_00851 has a product template (PT) domain and a N-terminal ACP transacylase similar to the one involved in aflatoxin biosynthesis (SAT).

### Differentially expressed β-lactamases play an unknown role in fungi

Five β-lactamase-related (IPR001466) genes are upregulated (1.7 < L2FC < 7.1) while one is downregulated (L2FC = -4.0). In bacteria, β-lactamases confer antibiotic resistance against β-lactams which are produced by fungi to target bacterial peptidoglycan, with few examples of fungicidal β-lactams (Arnoldi *et al*., 1990). Yet, there are few β-lactamase-related genes characterized in fungi so the computational annotation of β-lactamases in fungal genomes is likely too specific and does not account for potential diversity of lactamases (Gao *et al*., 2017).

## DISCUSSION

We used an RNA-seq experiment to understand the transcriptional responses of the EHB *Luteibacter* sp. 9143 and foliar endophyte *Pestalotiopsis* sp. 9143 when grown together in co-culture vs. axenically. Based on previous experiments and preliminary data, we predicted that the bacterium would be mildly parasitic on the fungus, and would respond to co-culture with its host by altering transcription of genes related to chemotaxis, motility, nutrient acquisition, and secretion systems. Similarly, for the fungus, we predicted changes related to carbohydrates such as cellulose and the upregulation of genes related to repair mechanisms and responses to infection related to symbiosis with *Luteibacter*. These predictions were upheld by our analyses.

In cultures used for the transcriptomic study, we observed a greater density of suspended cells of *Luteibacter* sp. 9143 in co-culture with *Pestalotiopsis* sp. 9143 compared to the axenic bacterial culture. This supports previous findings that the bacterium is not able to obtain sulfur in the form of sulfate when grown in pure culture, and that growing in co-culture with its host fungus can supplement this deficiency (Baltrus and Arnold, 2017). Upregulation of genes encoding ribosomal proteins during co-culture was consistent with the increased bacterial growth. At the same time, we observed a greater biomass of *Pestalotiopsis* sp. 9143 grown in pure culture compared to that of the fungus growing with its EHB in co-culture, implying that fungal growth is inhibited in the presence of *Luteibacter* sp. 9143. Whether or not this reduction in growth of the fungus in the presence of the bacterium influences interactions with host plants remains unknown, but should be explored further.

Given the reduction in growth by *Pestalotiopsis* sp. 9143 in co-culture, the fungus may be undergoing a defense response to *Luteibacter* sp. 9143. Indicators of this include the upregulation of diverse transporters and β-lactamase genes, which could be turned on in response to or anticipation of antifungal compounds. For example, a *Fusarium verticillioides* “metallo-β- lactamase” protein actually confers resistance to plant-derived, antifungal γ-lactams (Glenn *et al*., 2016). Similarly, the upregulated lactamase genes in *Pestalotiopsis* sp. 9143 may play a role in countering antifungal compounds or in improving the environment for putative bacterial partners. Furthermore, it is likely at least a subset of the upregulated transporters mediate resistance to antifungal compounds, especially four differentially expressed genes (KJ359_009703, KJ359_010943, KJ359_010406, and KJ359_003031) that have conserved domains from the pleiotropic drug resistance protein family (PF06422). Antifungal resistance-related gene expression may be in response to *Luteibacter* sp. 9143 specifically or bacteria generally, and may be fruitful avenues to explore for host specificity in bacterial-fungal interactions.

The response of *Luteibacter* to *Pestalotiopsis* in co-culture includes bacterial cells in multiple stages of interaction, including free-living, attached, entering, and endohyphal, complicating interpretation of the transcriptional response. This may contribute to why we see low L2FC for cell wall- and membrane-associated genes in the fungus, as the majority of bacterial cells are likely external to the fungus. Still, we see the response of *Luteibacter* sp. 9143 in co-culture with *Pestalotiopsis*sp. 9143 as representing an immediate metabolic response to changes in environmental conditions and the initiation of association with the fungal partner.

This includes the downregulation of methionine metabolism, chemotaxis, and motility, and the upregulation of sorbitol and mannitol transport, and T6SS (**Fig. 2, 3**). In particular, we speculate that the upregulation of the T6SS may help *Luteibacter* to initiate and establish symbiosis with *Pestalotiopsis*, although recognize that the bacterium may be using the T6SS for other purposes such as nutrient acquisition or even defense (Cianfanelli *et al*., 2016; Si *et al*., 2016; Bayer-Santos *et al*., 2019). Whereas it is difficult to understand which metabolites may be exchanged during this interaction from these genomic data, future studies using metabolomics may provide additional insight (e.g., Spraker *et al*., 2016).

Metabolic studies also could illuminate the role secondary metabolites play in facilitating the establishment of fungal-bacterial partnerships, including both recruitment and invasion. In the *R. microsporus* and *Mycetohabitans* interaction, secondary metabolites are not thought to play a large role, as only one NRPS is upregulated and one PKS downregulated in *R. microsporus* (i.e., the natural host strain) when in contact with the bacterium (Lastovetsky *et al*., 2020). In contrast the *Ralstonia solanacearum* lipopeptide, ralsolamycin, induced substantial developmental shifts in host fungi, enhancing bacterial entry into fungal chlamydospores (Spraker et al., 2016). In our analysis, *Pestalotiopsis* differentially expresses the core biosynthetic gene in 13 secondary metabolite clusters, but only Cluster 10.1 had all predicted biosynthetic genes differentially expressed. Given the two PKS genes in Cluster 10.1, it is possible that two separate metabolites are made, but the co-regulation and tight clustering support a single metabolic pathway. For example, a single *Aspergillus nidulans* cluster creates asperfuranone using two PKS genes, one highly reducing and the other not, along with five additional genes (Chiang et al., 2009). Cluster 10.1 does not appear to be related to the asperfuranone cluster based on the difference in accessory gene content and lack of homology between KJ359_00848 and KJ359_00851 and the asperfuranone PKS genes, though it does similarly contain one PKS predicted to be highly reducing and one not. Notably, this cluster is present in *Neopestalotiopsis clavispora*, *Neopestalotiopsis* sp. 37M, and *Pestalotiopsis microspora* at nearly 80% protein identity, suggesting conservation in closely related fungi. We also observed *Luteibacter* sp. 9143 to upregulate genes predicted to code for multidrug efflux pumps, which may be in response to the production of secondary metabolites by *Pestalotiopsis* sp. 9143 and assist the bacterium in colonizing its host, especially if the fungus is producing them as a defense response to bacterial invasion.

Our study supports a more direct exchange of primary metabolites, as *Luteibacter* sp. 9143 cannot utilize the sulfate in minimal media as a sulfur source, thus requiring methionine or other organic sulfur compounds generated by the fungus (Baltrus and Arnold, 2017).

Methionine is limited in the plant apoplast and methionine synthases are critical in plant pathogenic ascomycetes for survival *in planta* (Solomon *et al*., 2000; Saint-Macary *et al*., 2015). The small upregulation of methionine synthesis genes by the fungus may indicate a feedback response to having sulfur-containing compounds be depleted by the presence of *Luteibacter* sp. 9143, or may be more actively encouraged by the bacterium in some way. In *Saccharomyces cerevisiae*, excess methionine is tied to upregulation of synthesis of the cofactor pyridoxal-5’- phosphate (PLP) (Walvekar & Laxman, 2019), as part of an anabolic program leading to increased synthesis of amino acids. The downregulation of PLP synthases in *Pestalotiopsis* sp. 9143 could be tied to decreased methionine availability based on bacterial use, potentially leading to less amino acid metabolism and contributing to the observed growth restriction.

Interestingly, genes predicted to be associated with methionine metabolism were downregulated in *Luteibacter* sp. 9143 (**Fig. 2B**), which also supports the hypothesis that the bacterium is acquiring methionine from *Pestalotiopsis* sp. 9143.

Altogether, our transcriptomic analysis supports the phenotypic findings we have observed in the *Pestalotiopsis-Luteibacter* partnership, including a dependence on the fungal host as a sulfur source for *Luteibacter,* a change in extracellular carbohydrate metabolism by *Pestalotiopsis* when *Luteibacter* is present, and inverse growth impacts on the partners during co-culture. Together, these findings support a working model in which *Luteibacter* switches from a motile to sedentary lifestyle in association with the fungus, which can provide it with methionine as a sulfur source. Other metabolites such as leucine, mannitol, or sorbitol may also be exchanged. The specific metabolites and signaling pathways leading to the transcriptional changes remain to be investigated, but may involve the highly upregulated bacterial two-component system or T6SS and the fungal PKS secondary metabolite Cluster 10.1 or NmrA repressors. Based on the diversity of EHB-fungal relationships, additional work to probe the mechanisms underlying this and other partnerships will be critical for informing a broader model of bacterial-fungal interactions, and especially how they relate to ecology in the phyllosphere.

## MATERIALS AND METHODS

We obtained *Pestalotiopsis* sp. 9143 from a living culture collection at the Robert L. Gilbertson Mycological Herbarium, University of Arizona, Tucson (ARIZ). The fungus was isolated originally as an endophyte from healthy, asymptomatic foliage of *Platycladus orientalis* (Cupressaceae; Hoffman and Arnold, 2008). *Pestalotiopsis* sp. 9143 was naturally infected with the endohyphal bacterium *Luteibacter* sp. 9143 at the time of isolation, and maintained a consistent infection throughout growth in culture on various media and vouchering in sterile water. The bacterium was isolated successfully in culture and a rifampicin-resistant strain (9143) was generated by plating on LB amended with 50 µg/mL rifampicin (Arendt *et al*., 2017).

To generate cured *Pestalotiopsis* sp. 9143, conidia from sporulating, 21 day-old cultures were transferred to a new Petri plate (60-mm) containing 2% malt extract agar MEA amended with four antibiotics: tetracycline (10 µg/mL), ampicillin (100 µg/mL), ciprofloxacin (40 µg/mL) and kanamycin (50 µg/mL; MEA+TACK). Total genomic DNA extracted from fungal mycelium was used for polymerase chain reaction (PCR) to screen for presence or absence of *Luteibacter* sp. 9143 by amplifying the bacterial 16*S* ribosomal RNA (rRNA) gene following (Shaffer *et al*., 2016). Successful amplification was assessed by mixing PCR products with SYBR green and running on a 2% agarose gel in Tris-EDTA buffer (1X).

### Culture conditions prior to RNA extraction

After confirming the absence of *Luteibacter* sp. 9143 in *Pestalotiopsis* sp. 9143 growing on MEA+TACK, conidia were transferred to a new MEA plate. We then revived *Luteibacter* sp. 9143 by streaking from glycerol stock onto lysogeny broth (LB) agar (1% NaCl; i.e., LB-Miller). Both fungal and bacterial cultures were incubated at ambient laboratory conditions. After 21 days, for *Pestalotiopsis* sp. 9143, six *ca.* 2-mm^2^ squares were excised from the growing edge of the fungal colony with a sterile toothpick, and transferred to a sterile 125-mL flask containing 50 mL fresh liquid M9 + glucose (2%) + methionine (10 mM) (high-methionine minimal) medium. In parallel for *Luteibacter* sp. 9143, a sterile toothpick was used to transfer bacterial cells from a single colony to a test tube containing 5 mL fresh liquid high-methionine minimal medium. Fungal and bacterial cultures were then incubated at 27°C shaking at 200 rotations per minute (rpm).

After 10 days we processed cultures to produce one culture flask for each of the following treatments: (1) *Pestalotiopsis* sp. 9143 grown alone (i.e., axenic fungus); (2) *Luteibacter* sp. 9143 grown alone (i.e., axenic bacterium); and (3) *Pestalotiopsis* sp. 9143 and *Luteibacter* sp. 9143 grown together (i.e., fungus + bacterium co-culture). Fungal inoculum was created by transferring mycelia from the fungal culture flask to a sterile 50-mL stainless steel Sorvall omni mixer homogenizer chamber assembly (5.08-cm blade, PTFE bearings; OMNI, Kennesaw, GA, USA) containing 15 mL fresh liquid high-methionine minimal medium for homogenization: (20s on, 1 min off, 20s on). Bacterial inoculum was created by diluting the liquid culture to an optical density at 600 nanometers (nm) (OD_600_) of 0.1. Five milliliters of mycelial suspension and of bacterial suspension were added to flasks with fresh liquid high-methionine minimal medium to create an axenic fungal, an axenic bacterial, and a fungal-bacterial co-culture, each at a final volume of 50 mL.

After four days we split cultures to produce three biological replicates for each of the three treatments above. We first transferred the contents of each culture flask to a sterile 50-mL Falcon tube (Corning, NY, USA), centrifuged at 11 000 rpm for 20 min (5430R, Eppendorf, Hamburg, Germany) to pellet cells, removed media by pipetting, and re-suspended in fresh liquid M9 + glucose (2%) + methionine (100 µM) (low-methionine minimal) medium to bring the total volume of each tube to 25 mL. We then transferred 5 mL of each culture to four sterile 125-mL flask containing 45 mL fresh liquid low-methionine minimal medium and incubated all cultures at 27°C shaking at 200 rpm. Previous work indicates *Luteibacter* sp. 9143 cannot utilize sulfate as a sulfur source during laboratory growth, but is able to grow in media supplemented with cysteine, methionine or high levels of thiosulfate, as well as in co-culture with its host fungus *Pestalotiopsis* sp. 9143 (Baltrus and Arnold, 2017). Although we used a high methionine medium in order to allow for robust growth of *Luteibacter* sp. 9143 up to this point, we now reduce its concentration in hopes of inducing association of *Luteibacter* sp. 9143 with *Pestalotiopsis* sp. 9143 in co-culture, as the bacterium appears to acquire sulfur from the host fungus (Baltrus and Arnold, 2017).

### Bacterial density and total biomass of experimental cultures

After three days and immediately prior to RNA extraction, we quantified the density of free-living *Luteibacter* sp. 9143 (i.e., those cells not associated with fungal mycelium in co-culture) in all cultures by removing 50 µL of liquid culture, diluting 1:1M, spreading 50 µL onto the surface of a Petri plate (100-mm) containing LB agar, and incubating at 27°C for 48 h prior to counting colony forming units (CFU). Following plating, we recovered and lyophilized all remaining tissue in each culture by first transferring to a 50-mL Falcon tube and centrifuging at 11 000 rpm for 20 min to pellet cells. We avoided manipulating the co-cultured flasks such as to isolate only bacteria associated with fungal mycelia, in order to prevent transcriptional responses (e.g., such as to washing), and note that the population of *Luteibacter* sp. 9143 in co-cultured flasks includes free-living, externally-associated, and endohyphal cells. We next transferred each pellet to a distinct, sterile 1.7-mL microcentrifuge tube with the lid punctured using a sterile needle, and immediately submerged samples in liquid nitrogen until boiling stops. We then transferred sample tubes to a pre-cooled lyophilizer flask and lyophilized samples for 24 h prior to storing at -80°C.

### Extraction of RNA from experimental cultures

For axenic fungal- and co-cultured samples, we transferred lyophilized tissue to a pre-weighed, sterile 1.7-mL microcentrifuge tube, ground tissue using a sterile pestle, and obtained total biomass by weighing and subtracting the weight of the tube. For axenic bacterial samples, we transferred 1.5 mL suspended cells to a sterile 1.7-mL microcentrifuge tube, pelleted cells, removed excess media, and flash froze before lyophilizing cells. To extract RNA, for each sample we used either 0.05 g tissue (*ca.* 100 µL of ground tissue; axenic fungal- and co-cultured samples) or entire lyophilized cell pellets (axenic bacterial samples). For each sample, biomass was added to a sterile 1.7-mL microcentrifuge tube containing sterile zirconium oxide beads. We then added 1 mL TRIzol to each sample tube, macerated samples using a beadbeater run for 2 min (power level 10), incubated samples on ice for 20 min, and centrifuged at 12 000 rpm for 15 min at 4°C (5804 Eppendorf) to pellet tissue.

We next transferred supernatants of each sample to sterile 1.7-mL microcentrifuge tube, added 250 µL chloroform, and mixed all samples uniformly by placing in a single tube rack, shaking the rack for 15 s, allowing to rest for 15 s, and shaking again for 15 s. We then incubated samples at ambient laboratory conditions for 5 min, centrifuged at maximum speed for 15 min at 4°C (5804 Eppendorf) to separate phases, transferred 400 µL of the upper, aqueous layer of each sample to a distinct, sterile 1.7-mL microcentrifuge tube discarding the pellet, and added 750 µL of pre-chilled (-20°C) ethanol (100%) to each sample tube. We inverted sample tubes several times uniformly as above, and incubated them on ice for 10 min. Nucleotides were observed precipitating in each sample. We next centrifuged sample tubes at maximum speed for 15 min at 4°C as above, removed and discarded supernatant from each sample tube by pipetting, washed each pellet with 1 mL ethanol (75%), decanted the supernatant, and added 1 mL ethanol (75% with DEPC water) to each sample tube. We then centrifuged samples at 14 000 rpm for 5 min at 4°C, decanted the supernatant, and dried pellets by centrifuging in a vacuufuge at 12 000 rpm for 3 min at 30°C. We next re-suspended each pellet in 40 µL DEPC water and incubated sample tubes for 5 min at 65°C, prior to storage at -80°C.

We quantified the 260/280 (i.e., nucleotide/protein) and 260/230 (nucleotide/organic) ratios for each sample using a NanoDrop (pre-DNase ratios), prior to treatment with DNase using the DNA-free kit (Ambion AM1906, ThermoFisher, Waltham, MA, USA). Briefly, for each sample we transferred a volume equivalent to 20 µg total nucleotide to a sterile 1.7-mL microcentrifuge tube and added DEPC water to obtain a total volume of 45 µL. We then added DNaseI Buffer to each sample, vortexed gently, added 1 µL rDNase, vortexed gently, and incubated sample tubes for 30 min at 35°C. For all samples we then repeated adding 1 µL rDNase, vortexing, and incubating for 30 min at 35°C. We next added 5 µL DNaseI Inactivation Suspension to each sample tube and incubated at ambient laboratory conditions for 2 min, hand- vortexing every 30 s. We quantified 260/280 and 260/230 ratios for each sample again as above (post-DNase ratios), diluted each sample to a concentration of 400 ng/µL in DEPC water to produce a total volume of 50 µL, and submitted sample tubes to the University of Arizona Genetics Core (UAGC) for quality checking, rRNA depletion and poly-A selection, and sequencing.

### RNA sequencing

From all fungal RNA samples, fungal rRNA was depleted twice with the Ribominus Transcriptome Isolation Kit for yeast (Thermofisher) and poly-A selection to acquire fungal transcripts was done using the NEBNext Poly(A) mRNA magnetic isolation module (New England BioLabs). The remaining RNA from the co-cultured samples and the RNA from the axenic bacteria samples were depleted twice of bacterial rRNA using the Ribominus Transcriptome Isolation Kit for bacteria (Thermofisher). All subsequent quality checking was performed by the University of Arizona Genomics Core (UAGC) using an Agilent Bioanalyzer 2100 and RNA 6000 Pico Chips. Sequencing libraries were prepared using RNA TruSeq library construction kits (Illumina, San Diego, CA, USA) and sequenced on an Illumina HiSeq 2500 using TruSeq 2 x 100 base pair (bp) paired-end chemistry (Illumina) by UAGC.

Sequencing and assembly of *Pestalotiopsis* sp. 9143 genome

Genomic DNA of Pestalotiopsis sp. 9143 was prepared using following Arnold and Lutzoni (2007). Genomic DNA was sent to the Microbial Genome Sequencing Center (MiGS) for library preparation using an Illumina tagmentation kit and paired-end (2 x 150 bp) sequencing with a NextSeq 550 instrument as per Baym *et al*. (2015). Long-read sequencing was done in the Baltrus lab from the same genomic DNA sample. DNA was prepared for sequencing using the LSK109 ligation sequencing kit without shearing and using the Long Fragment Buffer. Reads were sequenced using an R9.4 flowcell in a MinION, with basecalling by Guppy (v.3.2.6).

Illumina and Nanopore sequencing reads were used to create a *de novo* hybrid assembly with MaSuRCA (v3.4.1) (Zimin *et al*., 2013; Zimin *et al*., 2017). The resulting assembled genome (**Table S5**) FASTA file was used as input for gene prediction and annotation by the funannotate (v1.7.4) pipeline (Palmer & Stajich, 2016). Repetitive contigs were removed and the remaining contigs sorted before repeat masking. RNA-seq reads assembled with Trinity (Haas *et al*., 2013) were used as transcript evidence for gene prediction of the softmasked genome.

Putative functional annotation by funannotate (**Table S6**) was informed by searches of UniProt DB version 2020_04, antiSMASH 5.0 (Blin *et al*., 2019), Phobius (Käll *et al*., 2007), InterproScan5 (Jones *et al*., 2014), dbCAN v9.0 (Yin *et al*., 2012), MEROPS v12.0 (Rawlings *et al*., 2013) and eggNOG-mapper v1.0.3 (Huerta-Cepas *et al*., 2017). Two scaffolds were flagged as part of the mitochondrial genome and annotated with MITOS (Bernt *et al*. 2013). The genome assembly and annotations were deposited in NCBI Genomes (JAHZSN000000000.1) and the raw reads were deposited in the Sequence Read Archive (SRR14629398; SRR14629399). All sequencing from this study can be accessed through BioProject PRJNA732082.

### Analysis of RNA-seq

Illumina data were quality controlled using QoRTs (Hartley and Mullikin, 2015). We mapped fungal reads onto the genome of *Pestalotiopsis* sp. 9143 (JAHZSN000000000.1), and bacterial reads onto that of *Luteibacter* sp. 9143 (JQNL00000000.1; Baltrus *et al*., 2017) using STAR v2.7.3 (Dobin *et al*., 2013). Mapped reads were subset with SAMtools (Li *et al*., 2009) and transcript abundances quantified using featureCount, part of the subread (v2.0.1) package (Liao *et al*., 2014) (**Tables S7, S8**). One axenic fungal sample had triple the reads from sequencing, so the reads were randomly downsampled with SAMtools to be more closely aligned with the number of reads from the other samples (**Table S8**). We carried out analysis of differential gene expression using the R package *DESeq2* (Love *et al*., 2014; R Core Team, 2021) with a false discovery rate (FDR) value of 0.01 as a threshold for differential expression. We looked for enrichment for specific gene ontology (GO) terms among all differentially expressed genes for both *Pestalotiopsis* sp. 9143 and *Luteibacter* sp. 9143 (*i.e.*, grown in co-culture vs. axenically). Separately for the fungus and the bacterium, we first generated a list of differentially expressed genes and corresponding GO annotations based on the reference genome, and then conducted a functional enrichment analysis using a Fisher’s Exact Test with a *p*-value of 0.01 (Conesa *et al*., 2005) using the R package *topGO* (Alexa and Rahnenfuhrer, 2021). Raw sequencing reads and processed featureCounts tables can be accessed through the NCBI Gene Expression Omnibus (GSE181155).

## AUTHOR CONTRIBUTIONS

AEA and DAB conceived of the study, with JPS and JES. AEA provided the original fungal and bacterial material. JPS and JES cultured the fungus and bacterium, cured the fungus of bacteria, and conducted experiments; JPS, JES, and MEC analyzed data; KH and DB conducted preliminary experiments crucial to the work; BS assisted with data analysis; MC assisted with genomic DNA extraction for *Pestalotiopsis*; JPS, JES, MEC, DB, and AEA led the development of the manuscript, with contributions from all authors.

## ACKNOWLEDGMENTS

An allocation of computer time from the UA Research Computing High Performance Computing (HPC) at the University of Arizona is gratefully acknowledged.

